# Screening of cell-virus, cell-cell, gene-gene cross-talks among kingdoms of life at single cell resolution

**DOI:** 10.1101/2021.08.13.456190

**Authors:** Dongsheng Chen, Zhihua Ou, Jiacheng Zhu, Peiwen Ding, Haoyu Wang, Lihua Luo, Xiangning Ding, Tianming Lan, Weiying Wu, Yuting Yuan, Wendi Wu, Jiaying Qiu, Yixin Zhu, Yi Jia, Yanan Wei, Qiuyu Qin, Runchu Li, Chengcheng Sun, Wandong Zhao, Zhiyuan Lv, Mingyi Pu, Shangchen Yang, Ashley Chang, Xiaofeng Wei, Fengzhen Chen, Tao Yang, Zhenyong Wei, Fan Yang, Yuejiao Li, Yan Hua, Huan Liu

## Abstract

The outbreak of severe acute respiratory syndrome coronavirus 2 (SARS-CoV-2) issued a significant and urgent threat to global health. The exact animal origin of SARS-CoV-2 remains obscure and understanding its host range is vital for preventing interspecies transmission. Previously, we have assessed the target cell profiles of SARS-CoV-2 in pets, livestock, poultry and wild animals. Herein, we expand this investigation to a wider range of animal species and viruses to provide a comprehensive source for large-scale screening of potential virus hosts. Single cell atlas for several mammalian species (alpaca, hamster, hedgehog, chinchilla etc.), as well as comparative atlas for lung, brain and peripheral blood mononuclear cells (PBMC) for various lineages of animals were constructed, from which we systemically analyzed the virus entry factors for 113 viruses over 20 species from mammalians, birds, reptiles, amphibians and invertebrates. Conserved cellular connectomes and regulomes were also identified, revealing the fundamental cell-cell and gene-gene cross-talks between these species. Overall, our study could help identify the potential host range and tissue tropism of SARS-CoV-2 and a diverse set of viruses and reveal the host-virus co-evolution footprints.

## Introduction

SARS-CoV-2 is a zoonotic virus that’s responsible for the coronavirus disease 2019 (COVID-19)^1,2^. Recent studies reveal that SARS-CoV-2 can infect various animals including bats, pangolins, cats, dogs, ferrets, minks, etc^3–9^ These findings were mainly based on epidemiological investigations or animal infection experiments. While these traditional methods are essential to elucidate *bona fide* viral infection in animals, it is impossible to carry out large-scale screening on the versatile species that might be susceptible to this pathogen, due to the unavailability of virus/animal/experimental resources. Host range assessment based on cellular receptor profiles may be an effective surrogate to narrow down the suspected host list. Different cellular surface proteins/ligands have been employed by distinct pathogens as their receptor to initiate attachment and cell entry, for example, angiotensin-converting enzyme 2 (ACE2) for SARS-CoV and SARS-CoV-2^3,10^, DPP4 for MERS-CoV^11^, sialic acid linked to galactose by α2,3/α2,6-linkages (Siaα2,3Gal/Siaα2,6Gal) for avian and human influenza viruses^12^, etc. Thus, understanding the associated receptor distribution may reveal the potential replicating niches of the viruses.

The COVID-19 pandemic has stimulated active investigations on the zoonotic origin of SARS-CoV-2 and this also urges us to expand this work to a bunch of other viruses that are related to respiratory, blood and encephalitis diseases. Identifying the host and tissue tropism is the first step towards understanding viral infection and pathogenesis, thus laying the foundation for the prevention and control of putative outbreaks in animals or humans. In this study, we constructed the single cell atlas for 20 species from livestock, poultry, pets and wild animals and surveyed the expression of over 100 receptors for 24 families at an unprecedented scale. The resulted resources are presented on a free online platform for free utilization by the academia and public community.

## Results

### Generation of single cell atlas for organs and PBMCs of 20 animals

In this study, we generated the single nuclei libraries for hedgehog (brain and kidney), chinchilla (brain and kidney), alpaca (frontal lobe, liver and lung) and hamster (frontal lobe, liver, kidney and heart) for 108,639 cells. The brain atlas mainly contained excitatory neurons (EX), inhibitory neurons (IN), microglia (MG), oligodendrocytes (OLG), oligodendrocyte progenitor cells (OPC), astrocytes (AST), endothelia cells (END). The kidney atlas is mainly composed of proximal tubule cells (PCT), loop of Henle cells (LOH), collecting duct principal cells (CD-PC), collecting duct intercalated cells (CD-IC), collecting duct transient cells (CD-trans), Podocytes (Podo), pericytes (PER), distal convoluted tubule cells (DCT), endothelial cells, smooth muscle cells (SMC), B cells. The lung atlas includes type □ alveolar cells (AT1), type II alveolar cells (AT2), ciliated cells (CC), endothelial cells, epithelial cells (EPI), fibroblasts (FIB). The heart atlas contains neuronal cells (NEU), cardiomyocytes, fibroblasts, mesothelial cells, macrophages, endothelial cells, pericytes. The liver atlas contains liver sinusoidal endothelial cells, Cholangiocytes, Kupffer cells, hepatocytes, hepatic stellate cells. (Figure 1a). The PBMC atlas for a total of 16 species (dalmatian pelican, black necked crane, red and green macaw, peacock, blue and yellow macaw, helmeted guineafowl, green cheeked parakeet, monk parakeet, sun conure, grey parakeet, snake, mongolian horse, tiger, alpaca, red necked wallaby, domestic guinea pig) was constructed, resulting in a total of 79,761 cells after QC (Figure 1b). The major immune cells, including T cells (T), B cells (B), macrophages (MAC), natural killer cells (NK), dendritic cells (DC) were identified based on the specific expression of canonical markers.

**Figure 1.**
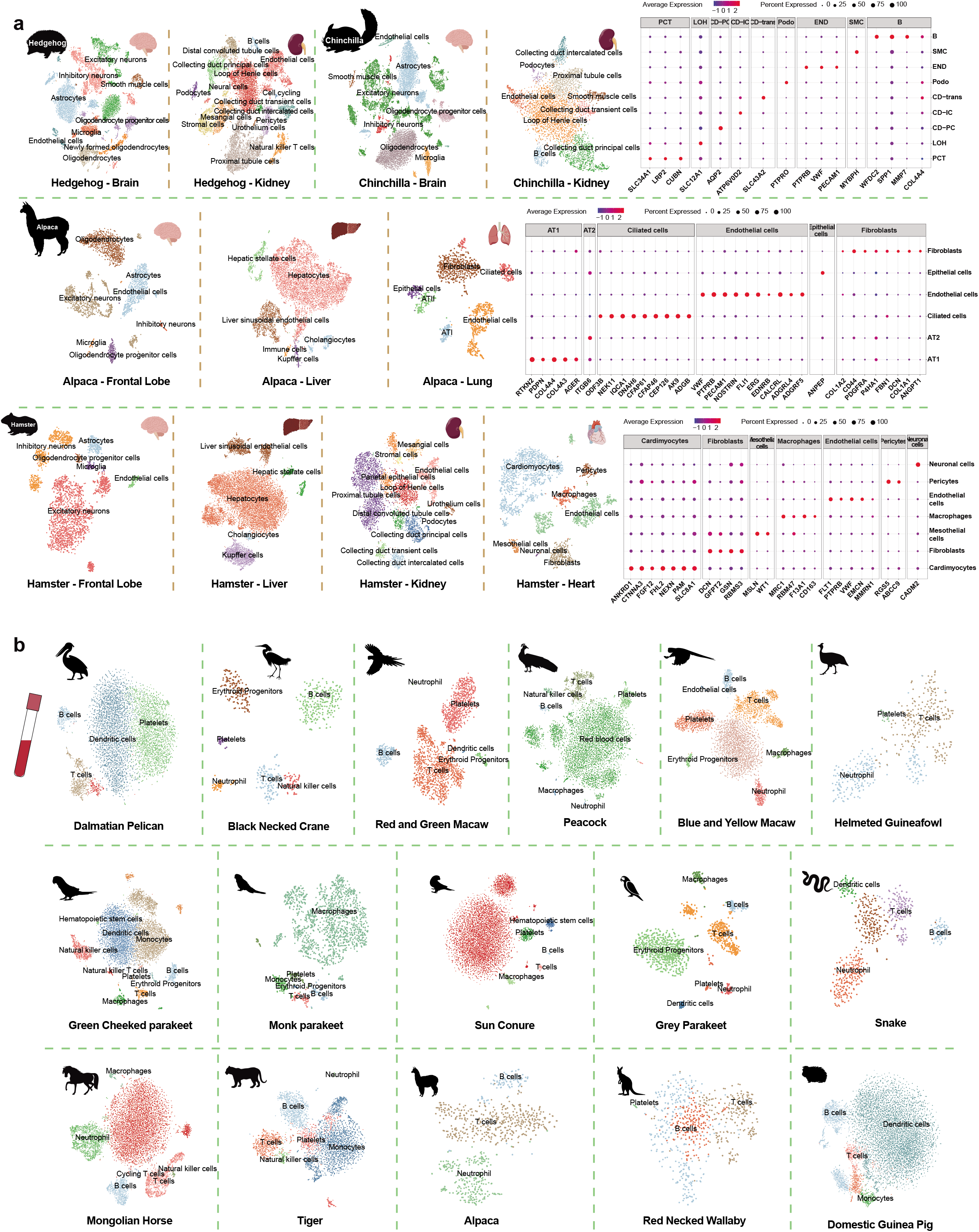
Generation of single cell atlas of hedgehog, chinchilla, alpaca, hamster and PBMC single cell atlas of 16 species. **a**. The single nuclei libraries for hedgehog (brain and kidney), chinchilla (brain and kidney), alpaca (frontal lobe, liver and lung) and hamster (frontal lobe, liver, kidney and heart) were generated with a total number of 108,639 cells. **b**. The PBMC atlas of 16 species including dalmatian pelican, black necked crane, red and green macaw, peacock, blue and yellow macaw, helmeted guineafowl, green cheeked parakeet, monk parakeet, sun conure, grey parakeet, snake, mongolian horse, tiger, alpaca, red necked wallaby, domestic guinea pig was constructed.

### Screening of SARS-CoV-2 entry factors and cofactors in the tissues of hedgehog, chinchilla, alpaca and hamster

Next, we screened the expression patterns of several SARS-CoV-2 entry factors and cofactors in tissues of hedgehog (kidney and brain), chinchilla (kidney and brain), alpaca (lung, liver and the frontal lobe) and hamster (liver, kidney, heart and the frontal lobe). Angiotensin-converting enzyme 2 (*ACE2*) and the tyrosine-protein kinase receptor UFO (*AXL*) are the cellular receptors responsible for SARS-CoV-2 infection^3,13^, while trans-membrane serine protease (*TMPRSS2*), Neuropilin-1 (*NRP1*) and the high-density lipoprotein scavenger receptor B type 1 (*SCARB1*) are cofactors promoting the ACE2-dependent entry of SARS-CoV-2^14–16^. Co-expressions of *ACE2* and its cofactors were identified for the four species (Figure 2, S1), with enrichment in the brain endothelial cells and kidney pericytes of hedgehog, in the brain astrocytes and kidney B cells of chinchilla, in the liver hepatocytes of alpaca, and in multiple cell types of hamster (the cardiomyocytes of heart, distal convoluted tubule cells, collecting duct transient cells and collecting duct intercalated cells of kidney, and liver sinusoidal endothelial cells, Kupffer cells and hepatocytes of liver). Intriguingly, *ACE2* was not detected in the lung of alpaca and the frontal lobe of hamster. *AXL* was detected in all tissues examined for the four species, especially in the microglia and endothelial cells in the frontal lobe of alpaca. In addition to *TMPRSS2*, we also screened for the expression of other members of the trans-membrane serine protease family, which displayed differential expression profiles in the organs of the four animals (Figure 2, S1).

**Figure 2.**
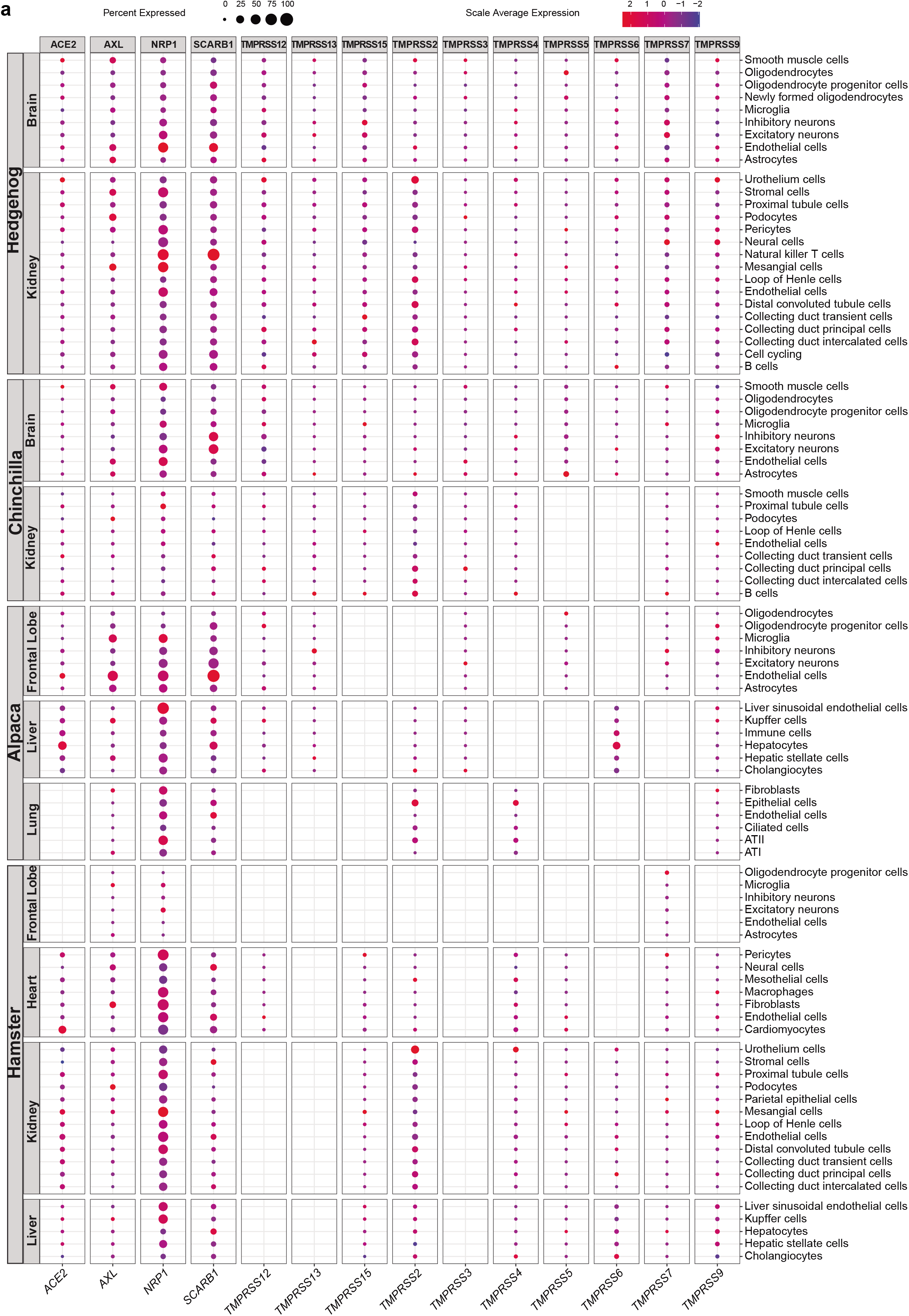
Screening of SARS-CoV-2 entry factors and cofactors in multiple species. The expression patterns of SARS-CoV-2 entry factors and cofactors, including angiotensin-converting enzyme 2 (*ACE2*), the tyrosine-protein kinase receptor UFO (*AXL*), Neuropilin-1 (*NRP1*), the high-density lipoprotein scavenger receptor B type 1 (*SCARB1*), trans-membrane serine protease (*TMPRSS2*) and the trans-membrane serine protease family, in tissues of hedgehog (kidney and brain), chinchilla (kidney and brain), alpaca (lung, liver and the frontal lobe) and hamster (liver, kidney, heart and the frontal lobe) were screened.

### Screening of viral receptors in PBMC data set of 23 species covering mammals, birds, reptiles and fish

Human immunodeficiency virus, feline immunodeficiency virus, Zika virus, Ebola virus, dengue fever virus, human herpes virus type 1, EBV virus, influenza virus, hepatitis G virus, enterovirus 71, hepatitis C virus and chikungunya mosquito virus was reported to infect human PBMC. We screened the expression of 33 associated viral entry factors (*CXCR4*, *TNFRSF4*, *CD4*, *CCR5*, *CLEC4M*, *CD209*, *AXL*, *TYRO3*, *NPC1*, *MERTK*, *CLEC4G*, *HAVCR1*, *CLDN1*, *RPSA*, *CLEC5A*, *ITGAV*, *ITGB6*, *ITGB8*, *NECTIN1*, *NECTIN2*, *TNFRSF14*, *EGFR*, *ANXA5*, *CACNA1C*, *UVRAG*, *HLA-DRA*, *CD74*, *HLA-DRB1*, *SCARB2*, *SELPLG*, *LDLR*, *CD81*, *PHB*) in the PBMC of 23 species, using the self-sequenced dataset for the above-annotated 16 species and the public dataset for 7 species including human, monkey, cat, rabbit, mouse, pigeon and zebrafish (Figure 3). ANXA5, ITGAV and NPC1 were commonly detected in the PBMC of the 23 in study. CXCR4, CD74, CD81 and SCARB2 were abundantly detected in mammals, birds, and reptiles, but was not found in zebrafish. *RPSA*, a receptor for adeno-associated virus, Sindbis virus, Venezuelan equine encephalitis virus, classical swine fever virus, dengue virus, was abundantly expressed in 20 out of the 23 species investigated including zebrafish.

**Figure 3.**
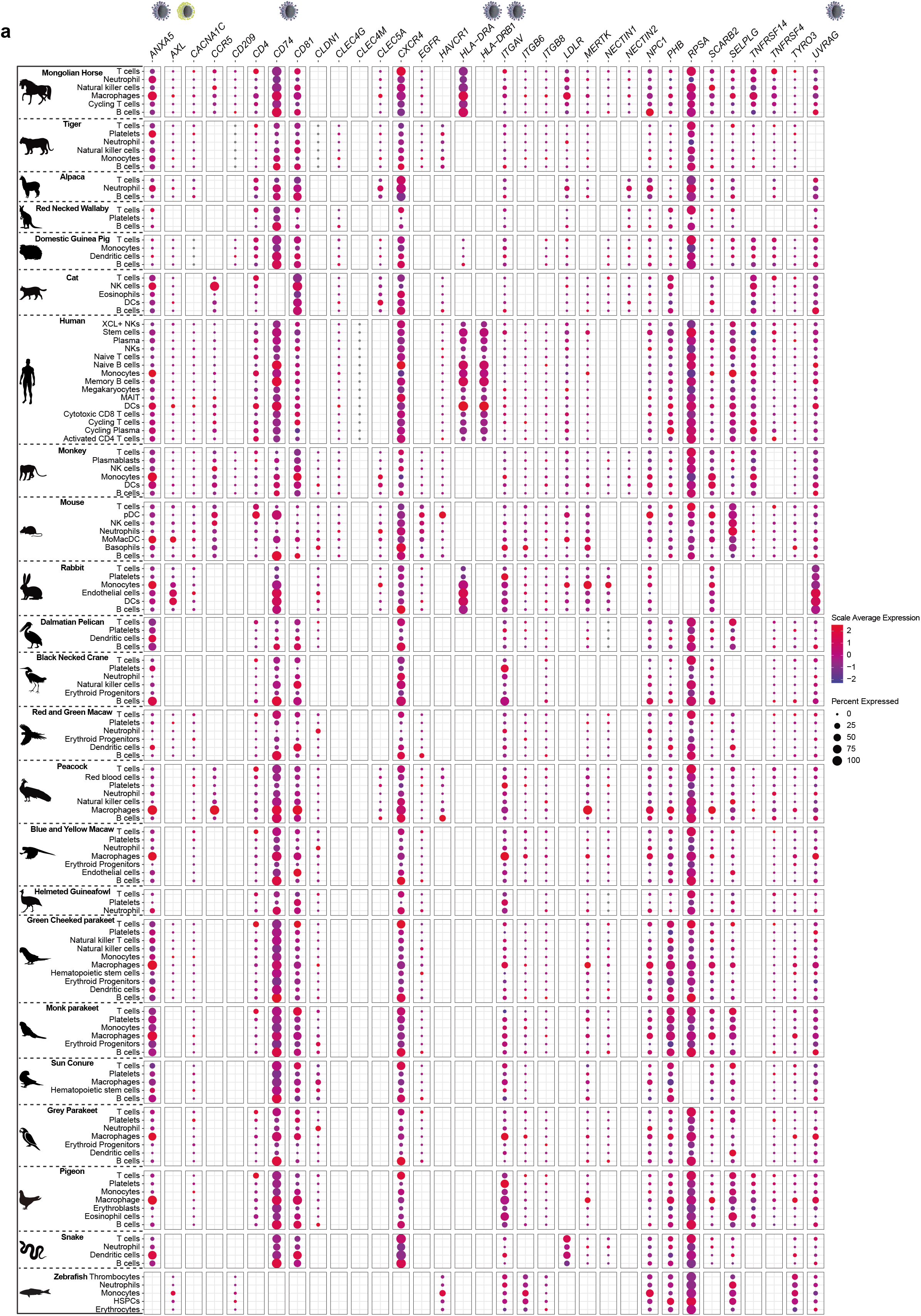
Cross-species screening of viral receptor expressions in the PBMC datasets. The expression of 33 associated viral entry factors in the PBMC of 23 species was screened, identifying some common and species-specific viral receptors.

### Screening of entry factors for neurotropic viruses in the brain tissues of mammals and reptiles

Although the brain is not the initial contact site of viral infection, neurological complications can be caused by neuroinvasive viruses including polioviruses, rabies viruses, flaviviruses, herpes simplex viruses, measles. In addition to the neuroinvasive viruses, a variety of other viruses such as coronaviruses and influenza viruses, are also capable of infecting the brain tissues under certain circumstances. To explore the putative target cells and hosts of the neurotropic viruses, we systematically screened the expression of 40 viral receptors in the brain tissues of 11 species covering 9 mammals (human, mouse, pig, alpaca, hamster, hedgehog, chinchilla, civet, mink) and 2 reptiles (lizard, turtle). *NCAM1*, *ITGB8*, *ITGAV*, *SCARB2*, *IDE*, *UVRAG*, ANXA5, CACNA1C and EGFR, were found to be widely expressed the mammals and reptiles studied (Figure 4).

**Figure 4.**
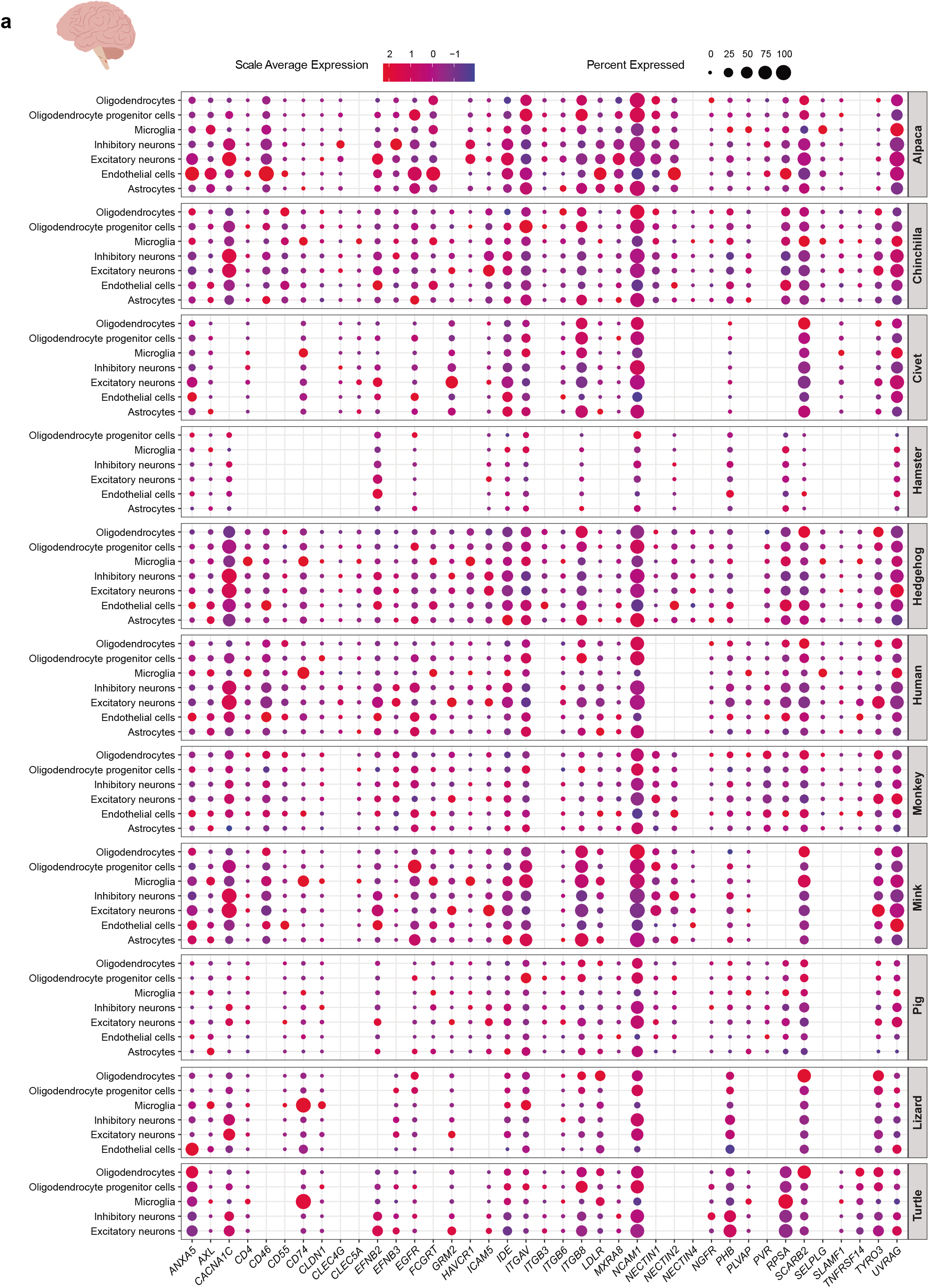
Cross-species screening of neurotropic virus target cells in brain tissues. The expression of 40 receptors for neurotropic viruses in the brain tissues of 11 species covering 9 mammals (human, mouse, pig, alpaca, hamster, hedgehog, chinchilla, civet, mink) and 2 reptiles (lizard, turtle) was screened.

### Cross-species screening of respiratory virus target cells in lung tissues

Respiratory syncytial viruses contribute to infections of lungs and respiratory tracts. Here, we selected 24 viral receptors corresponding to typical respiratory syncytial viruses and screened the expression patterns of their receptors (Figure S2a). Integrins are a family of transmembrane receptors which could facilitate both cell-cell and cell-extracellular matrix adhesion. Integrins are considered to commonly used viral receptors for a variety of nonenveloped and enveloped viruses (adenovirus, echovirus, hantavirus, foot-and-mouth disease virus, polio virus and other viruses). A total of 11 intergins (*ITGAV*, *ITGB6*, *ITGB8*, *ITGB1*, *ITGA4*, *ITGB3*, *ITGA3*, *ITGB5*, *ITGA6*, *ITGA2*, *ITGA5*) were reported to mediate viral entry. ITGA5, a receptor for human parvovirus B19, adeno-associated virus-2, human metapneumovirus, foot-and-mouth disease virus, was found to be highly expressed in endothelial cells of dog, bat and lizard (Figure S2b). UVRAG, receptor for influenza virus and vesicular stomatitis virus, was highly expressed in cat ATII and B cells, civet_fibroblasts, dog_macrophages (Figure S2c). RPSA, a receptor for a group of viruses, such as classical swine fever virus and dengue virus, was highly expressed in secretory cells of hamster (Figure S2d).

### Exploring tissue tropism and cell tropism of viruses at single cell resolution in human data set

Human alphaherpesvirus 1 (HHV-1) can infect human, pig and mouse via several receptors (ITGAV, ITGB6, ITGB8, NECTIN1, NECTIN2, TNFRSF14, CD209). HHV-1 infection was detected in human brain, oral mucosa, nose and conjunctiva and mouse mucocutaneous surfaces of skin, mouth and eyes while oral mucosa cell, tears and nasal mucosa cell were considered as target cells of HHV-1. Here, we checked the expression patterns of HHV-1 receptors over human brain, skin atlas. We observed the high expression of HHV-1 receptors in specific cell types. For example, ITGAV (AST, OLG, OPC), ITGB8 (AST, OLG, OPC, L2/3, L5/6 excitatory neurons), NECTIN1(PVRL1) (L2/3, L5/6 excitatory neurons, Neu-NRGN-1, IN), NECTIN2 (PVRL2) (endothelial, IN-SV2C, IN-VIP, IN-SST), TNFRSF14 (endothelial, L2/3 neurons), PVR, ITGB1 (IN, L5/6 neurons), ITGA4 (L2/3 neurons), IDE (L2/3, L5/6, L4, IN). Japanese encephalitis virus (JEV) could establish infection in human via microglia cells. In this data, we observed the specific expression of JEV receptor PLVAP in microglia cells. West Nile virus (WNV) infection causes endocytosis in human brain. Here, we detected the expression of ITGAV, an entry factor of WNV, in AST-PP, OPC, OLG, and endothelial. NCAM1, a receptor for Rabies lyssavirus, was widely expressed in every cell type of brain, except endothelial cells. Another Rabies lyssavirus, GRM2, was specifically expressed in L2/3, L5/6-CC, L4. *TYRO3*, receptor for Dengue virus, Zika virus, Marburg marburgvirus, Ebola virus, Lassa mammarenavirus, Lymphocytic choriomeningitis mammarenavirus, was specifically expressed in L2/3, L5/6-CC, L4, L5/6 and Neu-NRGN-1 of human brain.

In addition to central nervous system, we also checked the cell tropism of hepatitis virus in liver. SLC10A1 and GPC5, receptors for Hepatitis B virus, were specifically expressed in hepatocyte. Besides, SLC10A1 was enriched in hepatocyte of cat liver. Additionally, we checked the expression of viral receptors in digestive system. Human adenoviruses (Ads) are related to a broad range of infections, in particular upper respiratory and gastrointestinal tracts. CD46, CXADR were widely expressed in various cell types of colon (enterocyte, progenitor, stem cell, goblet, enteroendocrine, Paneth-like cell). CXADR was enriched in enterocyte of ileum. CD46, CXADR was widely expressed in different cell types of rectum.

Olfactory nerve function as a shortcut for virus entry into the CNS include influenza A virus, herpesviruses, poliovirus, paramyxoviruses, vesicular stomatitis virus, rabies virus, parainfluenza virus, adenoviruses, Japanese encephalitis virus. To test the putative target cells of those viruses, we systematically checked the expression of receptors (*ITGAV*, *ITGB6*, *ITGB8*, *NECTIN1*, *NECTIN2*, *TNFRSF14*, *CD209*, *NGFR*, *NCAM1*, *GRM2*, *EGFR*, *ANXA5*, *CACNA1C*, *CLEC4M*, *UVRAG*, *HLA-DRA*, *CD74*, *HLA-DRB1*, *CLEC4G*, *PLVAP*, *GKN3P*, *ITGB3*). Among which, receptor 1, 2, 3 were specifically expressed in respectively.

### Conservation of brain connectomes

In addition to cell-virus interactions, we explored the cross-talks between cells. To identify putative cellular communications, a ligand receptor-mediated interaction network was constructed for brain cells within 11 species. Overall, the source connectome topology of distinct species was quite similar, with human having the dominant source weight and hub scores (Fig. 5a). As for target analysis of network centrality, excitatory neurons cells function as signaling authority across all species. We next identified pan-conserved cellular connectivity, which may correspond to ancient signaling vectors inherited from common ancestors of mammals and fishes. In total, we detected 291 pairs of cell-cell interactions conserved among all 11 species, most of which were associated with the *NTRK2*.

**Figure 5.**
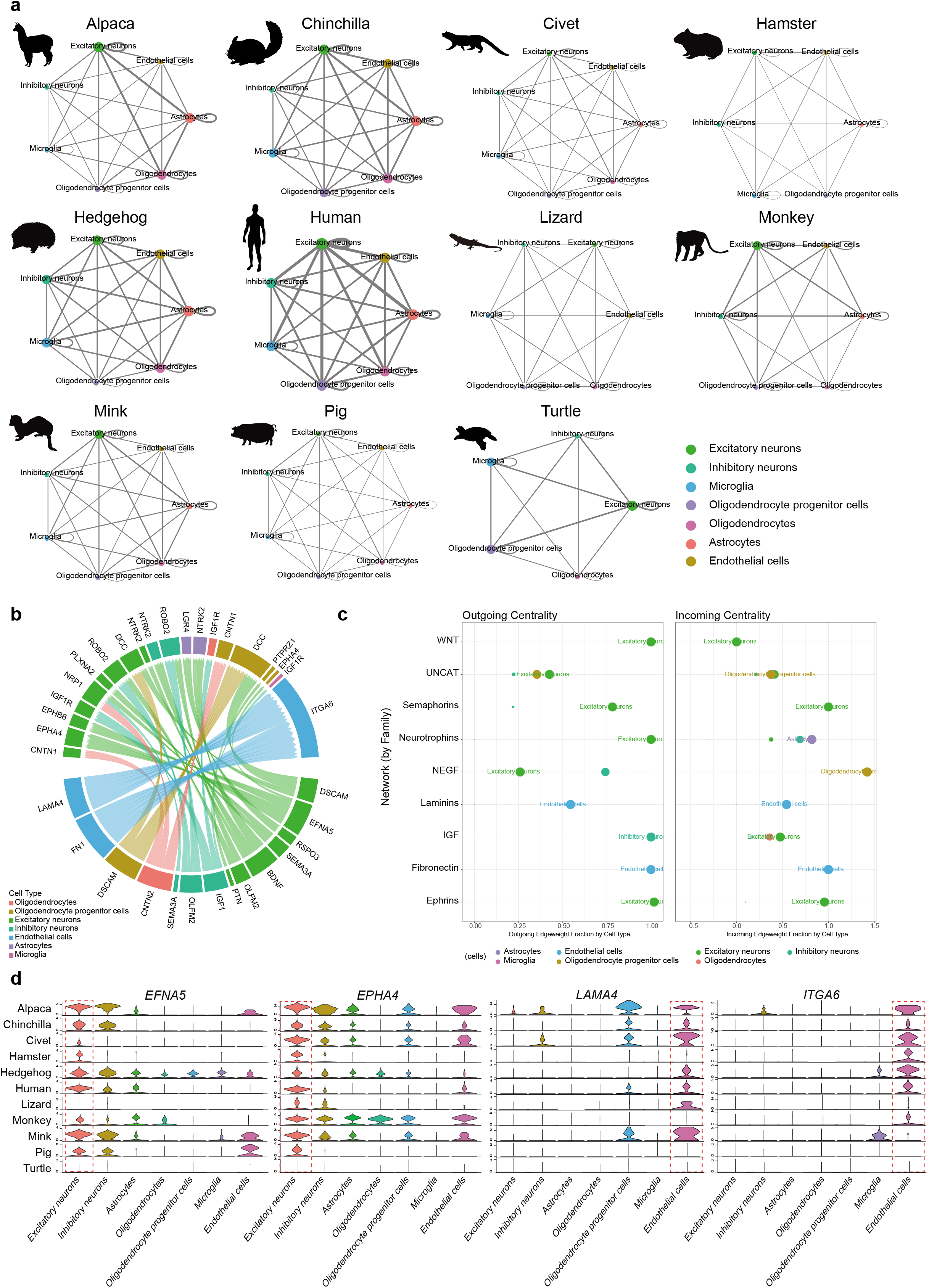
Conservation and divergence of brain cell connectomes. **a**. A ligand receptor-mediated interaction network was constructed for brain cells within 11 species. **b**. Analysis of pan-conserved cellular connectivity identified 291 pairs of cell-cell interactions conserved among all 11 species. **c**. The predominant cell types in cell-cell cross talk were identified as excitatory neurons cells, followed by endothelial cells. **d**. The stronger expression of EFNA5 (ephrin A5) and EphA4 (Eph receptor A4) in excitatory neurons, and LAMA4 (Laminin subunit alpha-4) and ITGA6 (Integrin alpha-6) in endothelial cells across species was demonstrated.

In addition to pan-conserved cellular cross-talk, we explored which cell types was predominant in cell-cell cross talk. In total, we found excitatory neurons cells appended in most families either outgoing centrality or incoming centrality and endothelial cells followed closely (Fig. 5c). For example, EFNA5 (ephrin A5) and EphA4 (Eph receptor A4) as the ligand and receptor of the ephrin family were commonly expressed in excitatory neurons cells. The ligand and receptor of laminins family, LAMA4 (Laminin subunit alpha-4) and ITGA6 (Integrin alpha-6), were also higher expressed in endothelial cells of most species (Fig. 5d). In summary, this study, for the first time, systematically revealed those highly conserved as well as lineage specific cell-to cell-signaling within vertebrate brain.

### Conservation of brain regulomes

To reveal the regulatory mechanisms underlying neural cells from the perspective of evolutionary biology, we predicted the regulomes in neural cells for all the 11 species, resulting in a total of 170,702 TF-target interactions in excitatory neurons cells and 383,544 TF-target interactions in inhibitory neurons cells (GENIE3 score > 0.01). The number of TF-target interactions conserved in at least 4 species, TF genes and target genes were expressed in 11 species ranged from 461 in excitatory neurons cells to 678 in inhibitory neurons cells (Fig. 6a, b). Encouragingly, several well-known regulators for excitatory neurons cells were active in the genetic regulatory network of the corresponding cell types, consistent with their expected regulatory functions. Of particular interest, we found a variety of regulatory circuits that were highly conserved among multiple species. Briefly, MEF2C (regulatory gene of excitatory neurons cells) with target genes (EPHA4, PEX5L, CNTN3 and PHACTR1), were conserved at least 8 species. In inhibitory neurons cells, the conserved regulatory relationships between MEF2C and other target genes (CACNA2D3, MMP16, PPARGC1A, RELN, SOX5, SOX6, ATRNL1 and KCNC2) were also associated at least 8 species. The regulatory functions of these transcription factors were inferred based on the enriched GO terms of their predicted target genes, the target genes of MEF2C were enriched in GO terms associated with positive regulation of neurogenesis, positive regulation of neurogenesis, positive regulation of neuron differentiation and regulation of neuron projection development either excitatory neurons cells and inhibitory neurons cells (Fig. 6c, d). We also found MEF2C had commonly expressed in all species (Fig. 6e). Overall, our study systematically identified conserved regulators for brain cells, including both well-recognized and novel regulators (Figure 6).

**Figure 6.**
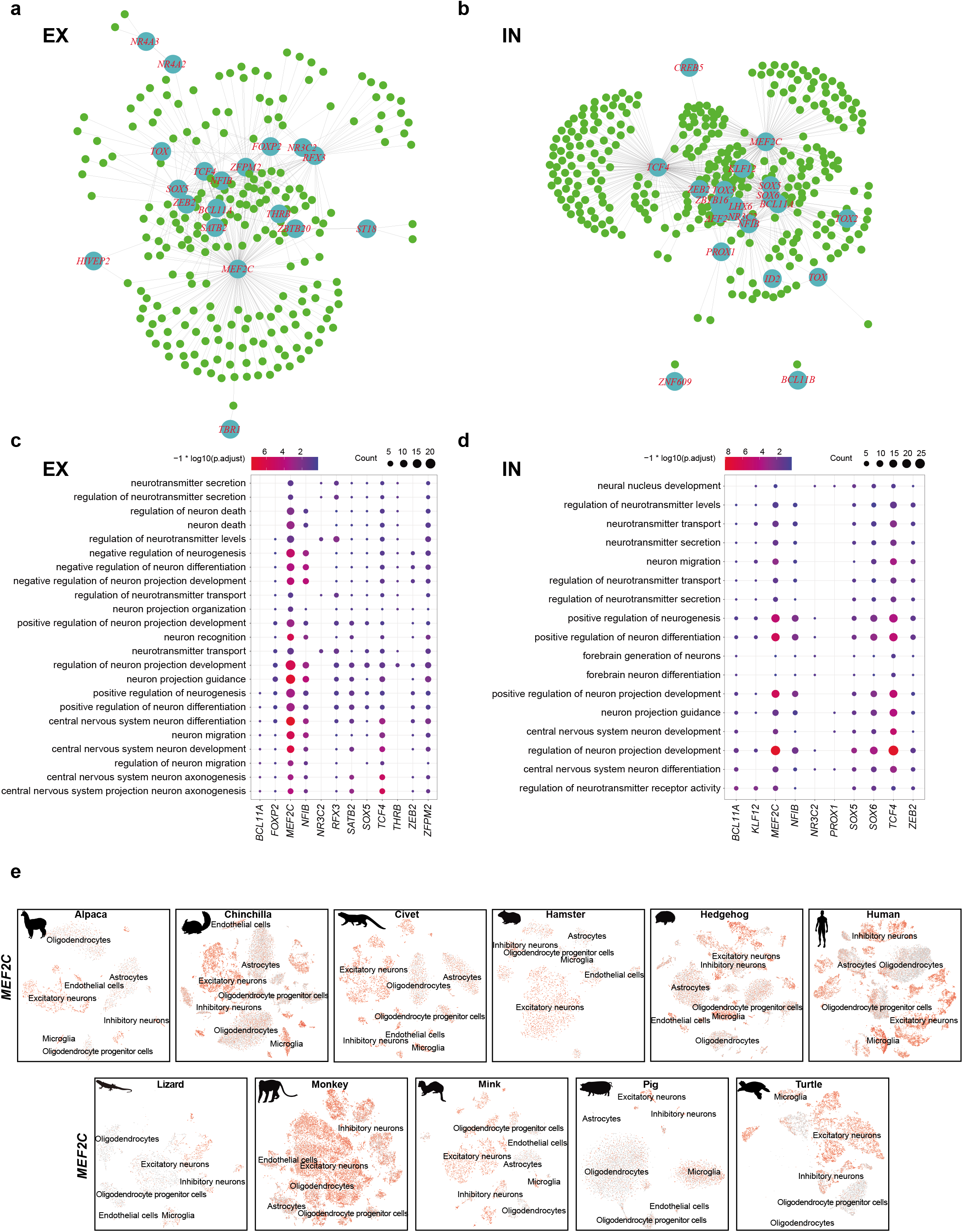
Conservation and divergence of brain cell regulomes. TF genes and target genes were expressed in 11 species ranged from 461 in excitatory neurons cells (**a**) to 678 in inhibitory neurons cells (**b**). The enrichment of the target genes of *MEF2C* in GO terms in excitatory neurons (**c**) and inhibitory neurons (**d**). The conserved expression of *MEF2C* in all species.

## Discussion

The widespread of SARS-CoV-2 makes it necessary to investigate the host and tissue tropism in animals. Here, we generated single-cell datasets from species including hedgehog (brain and kidney), chinchilla (brain and kidney), alpaca (frontal lobe, liver and lung) and hamster (frontal lobe, liver, kidney and heart) and screened the entry factors and cofactors of SARS-CoV-2 in the tissues of these species. Moreover, we screened the expression of receptors for other infecting viruses in PBMC sampled from more species covering mammals, birds, reptiles and fish. In addition, the entry factors for neurotropic viruses in the brain tissues of mammals and reptiles were investigated. These results revealed a comprehensive expression profile of virus receptors among species and may provide important evidence for understanding viral infection and pathogenesis. However, there are several limitations present in our study and the results should be interpreted cautiously. Firstly, we were only able to obtain scRNA data for tissues from 1-3 organs of each animal species in this study, due to difficulty in sampling and experiments. A more comprehensive characterization of the disease related organs may facilitate the interpretation of clinical symptoms induced by infections of various viruses in the animals. Secondly, the dataset of each organ for each species lacked biological nor technical replicates. Our results might be influenced by sampling bias. Thirdly, the expression profiles of the receptors were characterized based on the RNA sequencing data, not in their mature protein form with appropriate post-translational modifications. Some viruses, such as influenza A viruses, utilize sialic acid receptors and the sialyl linkage types to determine the differential binding specificity of avian and human adapted strains^12^. Although the transcriptome data could reveal the presence of the proteins associated with the receptors, the exact expression profiles of such kind of receptors could not be told by this data. Fourthly, we have only characterized the known receptor genes, other potential alternative receptors of the viruses were not included, but their distribution could also affect the tissue tropism of the viruses. Fifthly, both the expression profiles and the binding affinity between key cellular receptors and viral proteins are essential for viral entry into the host cells. Because protein products of the same ortholog may have different structures, and mutations in the binding domain may reduce or even abolish it binding ability to the corresponding viral surface proteins. The tissue tropism and host range of the viruses should be confirmed by experiments or epidemiological evidence.

In addition to investigate the cell-virus interactions, the datasets generated in this study can provide a comprehensive data source for the deep analysis of cross-talks between cells and genes. Here, we explored the cellular communications for brain cells within 11 species, and identified the excitatory neurons as the predominant cell type in cell-cell cross talk. The investigation of the regulomes in neural cells for all the 11 species identified conserved regulators for brain cells. These results suggest that the generated datasets can facilitate the research community for their personalized needs.

Overall, this study characterized the expression of receptors used by a variety of viruses in multiple tissues from mammals, birds, reptile and fish. The receptor profile may assist the exploration of tissue and host tropism of corresponding viruses, direct experimental design and guide the choice of appropriate animal infection models.

## Material and methods

### Single nuclei library construction and sequencing

Single nucleus libraries were constructed using Single Cell 3 □ GEM, Library & Gel Bead Kit v3 (PN-1000075) following the standard manufacturer’s instructions. Library conversion was performed using MGIEasy Universal DNA Library Preparation Reagent Kit to compatible with BGISEQ-500 sequencing platform.

### Single-nucleus RNA-sequencing data processing

Single-nucleus RNA-sequencing (snRNA-seq) data and gene expression matrix were obtained using Cell Ranger 3.0.2 (10X Genomics).

### Differential expression analysis

Differentially expressed genes (DEGs) were identified by “FindAllMarkers” function in Seurat. For DEGs of each cluster, we applied clusterProfiler Package for GO enrichment analysis.

### Expression of different virus receptors

All receptors of virus were collected from viral receptor database, Human Lung Cell Atlas (HLCA) database, or published articles. The expression of receptors in all cell types were displayed in dot plot using R package ggplot2.

### Public Data collection and processing

Public single cell data were downloaded from NCBI, GEO, Human Cell Atlas Portal, EBI single cell database.

## Acknowledgements

This work was supported by the China National GeneBank (CNGB). We thank the China National GeneBank for producing the sequencing data.

## Author Contributions

HL, XJ, YB and CDC conceived and designed the project. JCZ, WYW were responsible for sample collection and dissection. All authors participated in data interpretation, visualization and manuscript writing.

## Data availability

Single nucleus RNA-seq data that support the findings of this study have been deposited into CNGB Sequence Archive (CNSA) (Guo X, Chen F et al., 2020) of China National Gene Bank Data Base (CNGBdb) with accession code CNP0001889.

## Supplementary Figures

**Figure S1**. Screening of SARS-CoV-2 entry factors and cofactors in PBMC of multiple species.

**Figure S2**. Cross-species screening of respiratory virus target cells in lung tissues.

**Figure S3**. Exploring tissue tropism and cell tropism of viruses at single cell resolution in human data set.

## Reference

1. Zhou, P., Yang, X.-L., Wang, X.-G., Hu, B., Zhang, L., Zhang, W., Si, H.-R., Zhu, Y., Li, B. and Huang, C.-L. (2020) A pneumonia outbreak associated with a new coronavirus of probable bat origin. Nature, 579, 270–273.

2. Lam, T.T., Jia, N., Zhang, Y.W., Shum, M.H., Jiang, J.F., Zhu, H.C., Tong, Y.G., Shi, Y.X., Ni, X.B., Liao, Y.S. et al. (2020) Identifying SARS-CoV-2-related coronaviruses in Malayan pangolins. Nature, 583, 282–285.

3. Shi, J., Wen, Z., Zhong, G., Yang, H., Wang, C., Huang, B., Liu, R., He, X., Shuai, L. and Sun, Z. (2020) Susceptibility of ferrets, cats, dogs, and other domesticated animals to SARS–coronavirus 2. Science, 368, 1016–1020.

4. Munnink, B.B.O., Sikkema, R.S., Nieuwenhuijse, D.F., Molenaar, R.J., Munger, E., Molenkamp, R., Van Der Spek, A., Tolsma, P., Rietveld, A. and Brouwer, M. (2021) Transmission of SARS-CoV-2 on mink farms between humans and mink and back to humans. Science, 371, 172–177.

5. Sit, T.H.C., Brackman, C.J., Ip, S.M., Tam, K.W.S., Law, P.Y.T., To, E.M.W., Yu, V.Y.T., Sims, L.D., Tsang, D.N.C., Chu, D.K.W. et al. (2020) Infection of dogs with SARS-CoV-2. Nature, 586, 776–778.

6. Zhang, Q., Zhang, H., Huang, K., Yang, Y., Hui, X., Gao, J., He, X., Li, C., Gong, W., Zhang, Y. et al. (2020) SARS-CoV-2 neutralizing serum antibodies in cats: a serological investigation. bioRxiv, 2020.2004.2001.021196.

7. Halfmann, P.J., Hatta, M., Chiba, S., Maemura, T., Fan, S., Takeda, M., Kinoshita, N., Hattori, S.I., Sakai-Tagawa, Y., Iwatsuki-Horimoto, K. et al. (2020) Transmission of SARS-CoV-2 in Domestic Cats. N. Engl. J. Med., 383, 592–594.

8. Yuan, H., Yan, M., Zhang, G., Liu, W., Deng, C., Liao, G., Xu, L., Luo, T., Yan, H., Long, Z. et al. (2019) CancerSEA: a cancer single-cell state atlas. Nucleic Acids Res., 47, D900–D908.

9. Zhang, X., Lan, Y., Xu, J., Quan, F., Zhao, E., Deng, C., Luo, T., Xu, L., Liao, G., Yan, M. et al. (2019) CellMarker: a manually curated resource of cell markers in human and mouse. Nucleic Acids Res., 47, D721–D728.

10. Zhao, T., Lyu, S., Lu, G., Juan, L., Zeng, X., Wei, Z., Hao, J. and Peng, J. (2021) SC2disease: a manually curated database of single-cell transcriptome for human diseases. Nucleic Acids Res., 49, D1413–D1419.

11. Sun, D., Wang, J., Han, Y., Dong, X., Ge, J., Zheng, R., Shi, X., Wang, B., Li, Z., Ren, P. et al. (2021) TISCH: a comprehensive web resource enabling interactive single-cell transcriptome visualization of tumor microenvironment. Nucleic Acids Res., 49, D1420–D1430.

