## Supplementary figures and images for "Screening of cell-virus, cell-cell, gene-gene cross-talks among kingdoms of life at single cell resolution"

### Figure S1

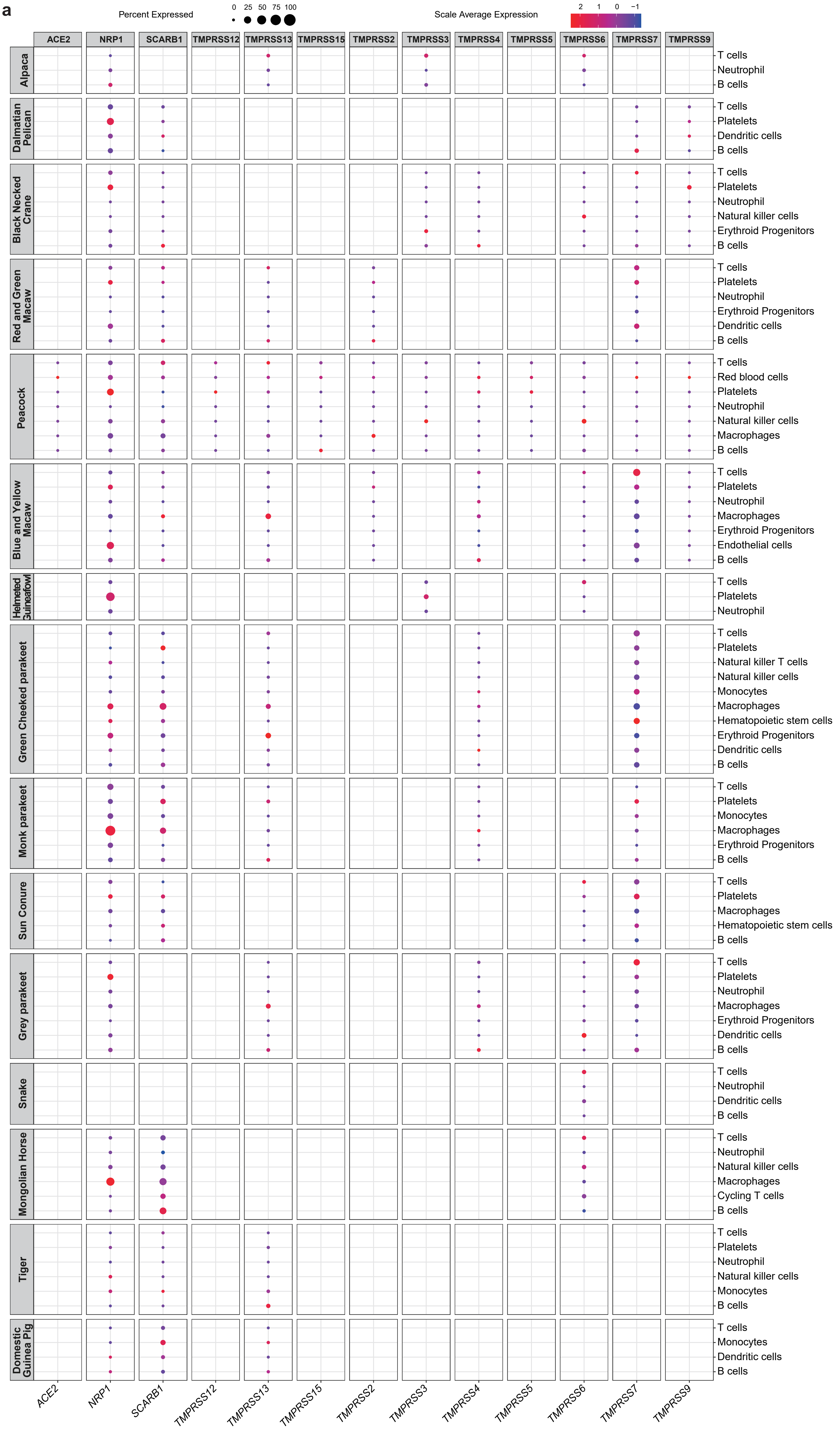

### Figure S2

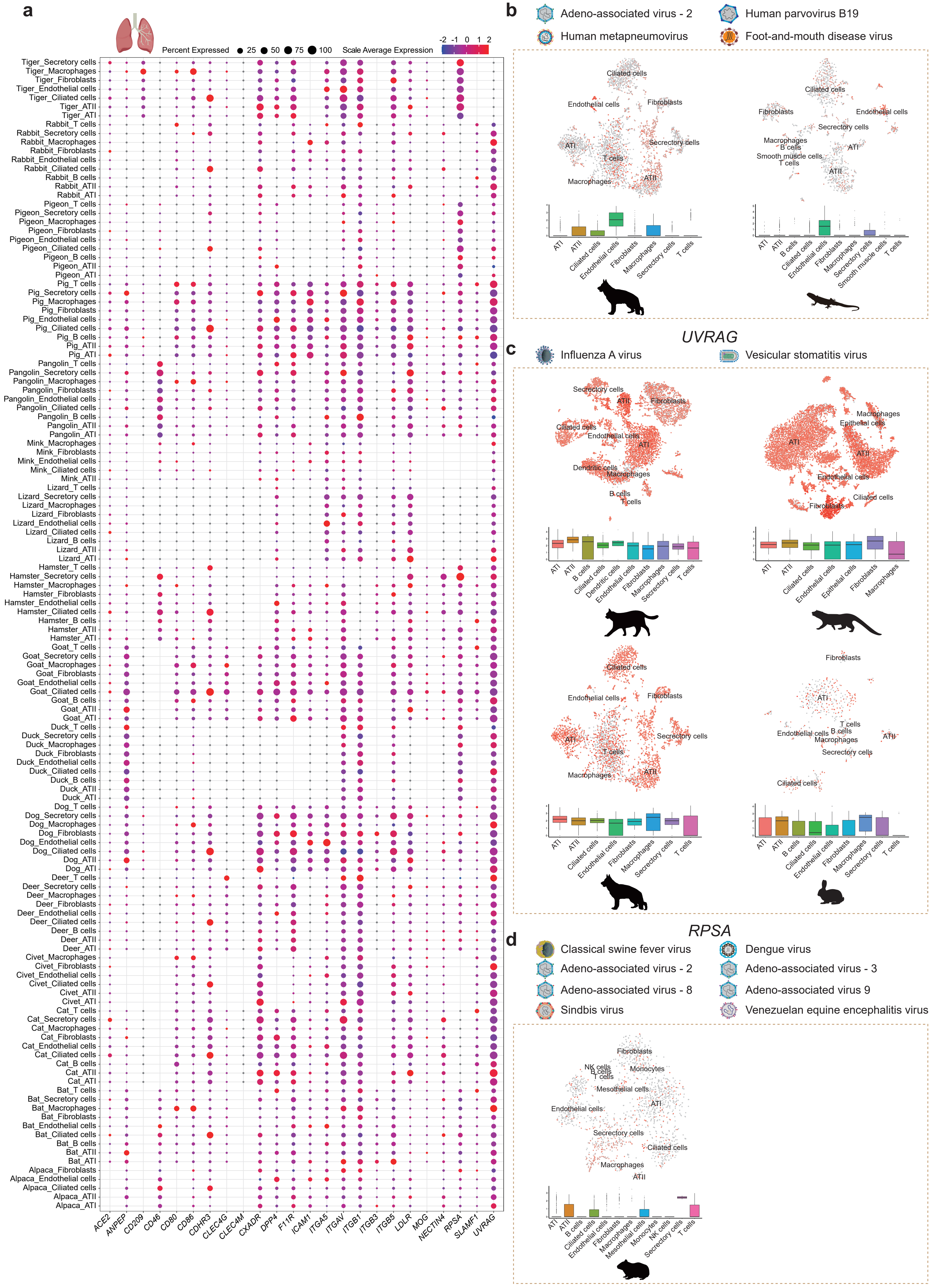

### Figure S3

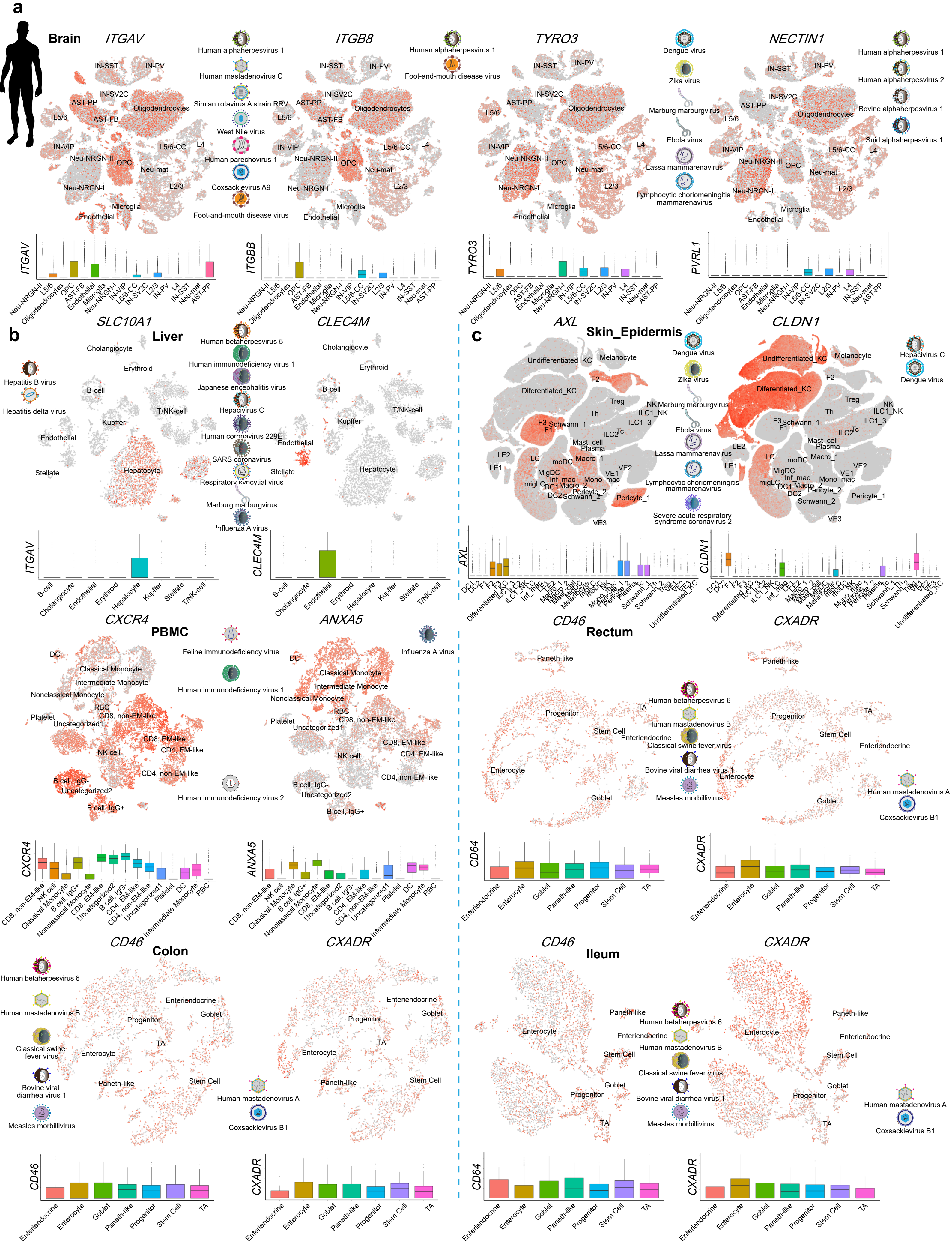
